# Alcohol use and cardiometabolic risk in the UK Biobank: a Mendelian randomization study

**DOI:** 10.1101/2020.11.10.376400

**Authors:** Joanna Lankester, Daniela Zanetti, Erik Ingelsson, Themistocles L. Assimes

## Abstract

Observational studies suggest alcohol use promotes the development of some adverse cardiometabolic traits but protects against others including outcomes related to coronary artery disease. We used Mendelian randomization to explore causal relationships between the degree of alcohol consumption and several cardiometabolic traits in the UK Biobank. We found carriers of the *ADH1B* Arg47His variant (rs1229984) reported a 26% lower amount of alcohol consumption compared to non-carriers. In our one-sample, two-stage least squares analyses of the UK Biobank using rs1229984 as an instrument, one additional drink/day was associated with statistically significant elevated level of systolic blood pressure (3.0 mmHg), body mass index (0.87 kg/m^2), waist circumference (1.3 cm), body fat percentage (1.7%), low-density lipoprotein levels in blood (0.16 mmol/L), and the risk of myocardial infarction (OR=1.50), stroke (OR=1.52), any cardiovascular disease (OR=1.43), and all-cause mortality (OR=1.41). Conversely, increasing use of alcohol was associated with reduced levels of triglycerides (−0.059 mmol/L) and HbA1C (−0.42 mmol/mol) in the blood, the latter possibly a consequence of a statistically elevated mean corpuscular volume among *ADH1B* Arg47His carriers. Stratifications by sex and smoking revealed a pattern of more harm of alcohol use among men compared to women, but no consistent difference by smoking status. Men had an increased risk of heart failure (OR = 1.76), atrial fibrillation (OR = 1.35), and type 2 diabetes (OR = 1.31) per additional drink/day. Using summary statistics from external datasets in 2-sample analyses for replication, we found causal associations between alcohol and obesity, stroke, ischemic stroke, and type 2 diabetes. Our results are consistent with an overall harmful effect of alcohol on cardiometabolic health at all levels of use and suggest that even moderate alcohol use should not be promoted as a part of a healthy diet and lifestyle.

## Introduction

The relationship between alcohol and cardiovascular disease is important to understand given the high prevalence of alcohol consumption [1,2]. In decades of epidemiological work, alcohol consumption has shown an inverse or J-shape association with multiple traits related to cardiometabolic health including Type 2 diabetes [3, 4], non-fatal and fatal coronary heart disease [5–7], ischemic stroke [8–10], atrial-fibrillation [11], and congestive heart failure [12].

One interpretation of these relationships has been that moderate drinking is beneficial to cardiometabolic health. A problem with this interpretation is that the relationship is inconsistent with that observed for some known risk factors; for example, alcohol has been directly associated with outcomes such as hypertension irrespective of the degree of intake [13]. A longstanding hypothesis to reconcile these observations stipulates that the negative effects on blood pressure are modest and are surpassed by the positive effects on HDL levels [14, 15] that are either directly affected by alcohol and/or a consequence of an improvement in insulin sensitivity [16]. However, this hypothesis has been challenged in the last decade by multiple randomized control trials of HDL-raising drugs, as well as Mendelian randomization studies that have failed to demonstrate the benefits on risk of cardiovascular disease (CVD) of pharmacologically or genetically raised HDL [17–19].

The extent to which observational studies can shed light on the relationship between alcohol and CVD is questionable due to confounding and reverse causality [20]. Alcohol use is related to cultural, socioeconomic, and lifestyle factors which cannot be fully accounted for in observational analyses. Furthermore, several studies have suggested substantial differences in the effects of alcohol on cardiometabolic traits between men and women [3, 4, 11, 12, 16, 21]. Mendelian randomization (MR) facilitates a comparison of groups of subjects that consume more vs. less alcohol that is free of confounding, allowing us to better understand the causal effect of consumption. In this study, we determine the causal relationship between alcohol and cardiovascular risk factors and disease in the UK Biobank by performing an instrumental variable analysis using a genetic variant in a gene *(ADH1B)* that is known to be responsible for the metabolism of alcohol and associated with the amount of alcohol consumed. We attempt to validate findings using summary statistics from external consortial studies for related phenotypes. Within the UK Biobank, we also take advantage of the large numbers to explore strength of associations stratified by sex and smoking status.

## Methods

### Study cohort

The UK Biobank is a prospective study of over 500,000 participants recruited in 2006–2010 [22]. Data collected from the participants included questionnaires, physical measures, sample assays, genotyping, and ongoing longitudinal hospital records. Participants were enrolled at age 40-69. This research has been conducted using the UK Biobank Resource under Application Number 13721. The Research Ethics Committee reference for UK Biobank is 16/NW/0274. The Stanford IRB reviewed the protocol and determined the research did not include human subjects as defined in 45 CFR 46, nor 21 CFR 56.

### Outcomes and quantitative traits

We extracted systolic and diastolic blood pressure, BMI, waist circumference, and body fat percentage from survey data which included physical measurement at the baseline clinic visit. We obtained lipids, blood count variables, and HbA1C from the biomarkers data. We extracted primary and secondary diagnosis disease outcomes from hospital data for myocardial infarction, stroke (hemorrhagic, ischemic, and any stroke), heart failure, atrial fibrillation, and a composite outcome of all cardiovascular events combined, as well as death from each of these disease outcomes according to the relevant ICD codes (S1.1). We derived type 2 diabetes status from a combination of diabetes-related questions and self-reported medications (S1.2).

### Main exposure and covariates

Our main exposure variable of interest was self-reported alcohol use by number of drinks per week or month and type of drink obtained from the survey data. Use of all types of alcohol was aggregated into total grams of alcohol intake per year [23], which was then transformed into equivalent daily glasses of wine (0, >0-1, >1-2, >2-3, and >3) to facilitate interpretability. Covariates also from survey data included sex, age, region of recruitment, socioeconomic status, ethnicity, smoking status, blood pressure medications, cholesterol-lowering medications, insulin and other diabetes drugs, and fasting status (for biomarkers) (S2).

We used Arg47His (rs1229984) in *ADH1B* as our genetic instrument for alcohol use. This variant is arguably the strongest and most established genetic predictor of self-reported alcohol use in European populations with a frequency of about 0.5% (Northern Europe) to 4% (Southern Europe) [24, 25]. The variant was directly genotyped using the UK Biobank array and thus no imputation of this variant was necessary.

### Statistical analysis

We characterized the observational relationship between alcohol use and continuous variable risk factors using linear regression for quantitative traits, logistic regression for type 2 diabetes, and Cox proportional hazards regression for cardiovascular events or death. The reference group for all observational analyses was current non-drinkers. For our Cox analyses, we defined the start of follow up as time of enrollment into the UK Biobank study and excluded those with a cardiovascular event prior to the questionnaire to minimize survivor bias. We created two models for each outcome in our observational analysis, one model minimally adjusted for typical epidemiologic covariates (sex, age, region of recruitment, socioeconomic status, ethnicity, and smoking status; for biomarker outcomes, fasting status was also included) and one model additionally adjusted for heart disease risk factors as well as medications that affect those risk factors (BMI and waist circumference, in all models other than those for anthropometric measures; SBP and DBP, in all models other than those for blood pressure; HbA1C and diabetic medications, in models other than for type 2 diabetes; LDL, HDL, and triglycerides, in all models other than those for lipids; and lipid-lowering, or anti-hypertensive medications).

We included all non-related individuals of European descent (for sample independence and to avoid population stratification) in the UK Biobank for our MR analyses. We grouped heterozygous and homozygous carriers (dominant model). We quantified the strength of the instrument variable using ANOVA and tested the relationship between the instrument variable and alcohol consumption using linear regression. We performed a one-sample MR using the two-stage least squares method with individual level UK Biobank data to estimate the causal effect on traits of consuming one additional drink per day on average. Given possible differences in drinking patterns by sex and smoking status, we also conducted stratified analyses in male, female, current smoker, and never smoker subgroups. We used the Breslow-Day test for heterogeneity of odds ratios to test whether outcomes varying substantially between strata were significantly different. For relevant outcomes, we conducted sensitivity analyses in subgroups not taking medications for diabetes, hypertension, and lipids. Lastly, we conducted an analysis of hemoglobin, hematocrit, and mean corpuscular volume levels by instrument variable status as an additional check on the potential confounding effects of anemia on our association analyses involving HbA1C. For replication in other datasets, we collected summary statistics for the largest available dataset of European or mostly European ancestry for each outcome (Table S3). We used these summary statistics for a two-sample MR, calculating the ratio of each summary statistic coefficient to the exposure-instrument variable coefficient from the UK Biobank. Analyses were done in R 3.6.3. Results were validated with the MendelianRandomization package.

## Results

UK Biobank participant characteristics are described in Table 1 and S5.1. At recruitment, 92% of participants consumed alcohol. About half of participants reported drinking alcohol 1-4 times per week, and an additional one fifth of participants reported drinking daily. On average, participants reported drinking 7.7 (± 9.4) glasses per week.

**Table 1.** Summary of characteristics of UK Biobank participants in datasets used for analyses, n (%) or mean(standard deviation)

| Variable | Total dataset | Observational analysis | Mendelian randomization |
| --- | --- | --- | --- |
| n | 502536 | 481150 | 337484 |
| Sex |  |  |  |
| Female | 273402 (54.4) | 267459 (55.6) | 181236 (53.7) |
| Male | 229134 (45.6) | 213691 (44.4) | 156248 (46.3) |
| Age (years) | 67.3 (8.1) | 67 (8.1) | 67.6 (8) |
| Region |  |  |  |
| England | 445883 (88.7) | 427194 (88.8) | 297645 (88.2) |
| Wales | 20808 (4.1) | 19936 (4.1) | 14824 (4.4) |
| Scotland | 35845 (7.1) | 34020 (7.1) | 25015 (7.4) |
| Townsend index | -1.3 (3.1) | -1.3 (3.1) | -1.6 (2.9) |
| Ethnic group |  |  |  |
| White | 472725 (94.1) | 452534 (94.1) | 337484 (100) |
| Asian or Asian British | 11456 (2.3) | 10847 (2.3) | 0 (0) |
| Black or Black British | 8061 (1.6) | 7863 (1.6) | 0 (0) |
| Mixed or other | 7517 (1.5) | 7279 (1.5) | 0 (0) |
| Don't know/refused | 2777 (0.6) | 2627 (0.5) | 0 (0) |
| Smoking status |  |  |  |
| Never-smoker | 273537 (54.4) | 265259 (55.1) | 183826 (54.5) |
| Current smoker | 52979 (10.5) | 50453 (10.5) | 33977 (10.1) |
| Former smoker | 173070 (34.4) | 162687 (33.8) | 118505 (35.1) |
| No response | 2950 (0.6) | 2751 (0.6) | 1176 (0.3) |
| Drinking status |  |  |  |
| Never-drinker | 22388 (4.5) | 21234 (4.4) | 10392 (3.1) |
| Current drinker | 460386 (91.6) | 441728 (91.8) | 315257 (93.4) |
| Former drinker | 18108 (3.6) | 16631 (3.5) | 11543 (3.4) |
| No response | 1654 (0.3) | 1557 (0.3) | 292 (0.1) |
| Drinking frequency |  |  |  |
| Daily or almost daily | 101774 (20.3) | 97438 (20.3) | 72270 (21.4) |
| Three or four times a week | 115445 (23) | 111123 (23.1) | 81462 (24.1) |
| Once or twice a week | 129297 (25.7) | 124157 (25.8) | 88747 (26.3) |
| One to three times a month | 55858 (11.1) | 53724 (11.2) | 37367 (11.1) |
| Special occasions only | 58012 (11.5) | 55286 (11.5) | 35411 (10.5) |
| Never | 40648 (8.1) | 38009 (7.9) | 21991 (6.5) |
| No response | 1502 (0.3) | 1413 (0.3) | 236 (0.1) |
| Alcohol (weekly equivalent glasses of wine) | 7.7 (9.4) | 7.7 (9.4) | 8.1 (9.5) |
| Systolic blood pressure (mmHg) | 139.7 (19.7) | 139.7 (19.7) | 140.2 (19.7) |
| Diastolic blood pressure (mmHg) | 82.2 (10.7) | 82.3 (10.7) | 82.2 (10.7) |
| Body mass index (kg/m<sup>2</sup>) | 27.4 (4.8) | 27.4 (4.8) | 27.4 (4.7) |
| Waist circumference (cm) | 90.3 (13.5) | 90 (13.4) | 90.3 (13.5) |
| Body fat percentage | 31.5 (8.5) | 31.5 (8.6) | 31.4 (8.5) |
| Cholesterol (mmol/L) | 5.7 (1.1) | 5.7 (1.1) | 5.7 (1.1) |
| LDL (mmol/L) | 3.6 (0.9) | 3.6 (0.9) | 3.6 (0.9) |
| HDL (mmol/L) | 1.4 (0.4) | 1.5 (0.4) | 1.5 (0.4) |
| Triglycerides (mmol/L) | 1.7 (1) | 1.7 (1) | 1.8 (1) |
| HbA1c (mmol/mol) | 36.1 (6.8) | 36 (6.5) | 36 (6.5) |
| Glucose (mmol/L) | 5.1 (1.2) | 5.1 (1.2) | 5.1 (1.2) |
| Type 2 diabetes | 25217 (5) | 22083 (4.6) | 15493 (4.6) |
| Coronary heart disease | 24047 (4.8) | 11297 (2.3) | 16102 (4.8) |
| All stroke | 11785 (2.3) | 6857 (1.4) | 8044 (2.4) |
| Ischemic stroke | 5081 (1) | 3217 (0.7) | 3427 (1) |
| Hemorrhagic stroke | 2814 (0.6) | 1608 (0.3) | 1877 (0.6) |
| Heart failure | 8956 (1.8) | 5128 (1.1) | 5921 (1.8) |
| Atrial fibrillation | 22197 (4.4) | 14292 (3) | 15483 (4.6) |
| Any cardiovascular disease | 52219 (10.4) | 30833 (6.4) | 35499 (10.5) |
| All-cause death | 20284 (4) | 17635 (3.7) | 13700 (4.1) |
| ADH1B status |  |  |  |
| Wildtype | 458807 (91.3) | 439084 (91.3) | 322519 (95.6) |
| Carrier | 26534 (5.3) | 25593 (5.3) | 14777 (4.4) |
| Homozygous minor allele | 1979 (0.4) | 1928 (0.4) | 188 (0.1) |

Exclusion of those with prior CVD (n= 21,386) yielded a dataset of 481,150 (Table 1). For our observational analysis (Figure S4.1-4S.2, Table S4.3-S4.6), increased alcohol use was directly related to higher systolic and diastolic blood pressure, total cholesterol, HDL, and atrial fibrillation in the fully-adjusted model. We observed a J-shape association (compared with no drinking, lower coefficient/odds ratio/hazard ratio with moderate drinking, but higher with heavy drinking) with BMI, waist circumference, body fat percentage, stroke (total, ischemic, and hemorrhagic), and all-cause death. Triglycerides, type 2 diabetes, myocardial infarction, heart failure and total cardiovascular disease also had a J-shape association, but with lower betas/odds ratio/hazard ratio at all drinking levels compared with non-drinkers. Alcohol was inversely associated with HbA1C. No pattern was observed between alcohol and LDL.

The Arg47His *ADH1B* variant was found in 4.4% of individuals in the UK Biobank, with only 188 homozygous for the variant. Our MR analysis (n=337,484) showed that those with the wildtype consumed 2.1 drinks/week more than carriers of the Arg47His variant in *ADH1B* (7,127 g/year or 8.2 glasses/week for wildtype vs 5,276 g/year or 6.1 glasses/week for carriers; F = 718) (Table 2). The two-stage least squares analysis showed that alcohol intake equivalent to one additional glass of wine per day was causally associated with higher systolic blood pressure (2.99 mmHg), BMI (0.87 kg/m^2), waist circumference (1.34 cm), body fat percentage (1.68%), cholesterol (0.160 mmol/L), LDL (0.155 mmol/L). Triglycerides (−0.059 mmol/L) and HbA1C (−0.420 mmol/mol) were significantly lower while diastolic blood pressure and HDL showed no difference. For disease events, one additional drink/week corresponded to higher risk of MI (OR = 1.50), all stroke (OR = 1.52), any cardiovascular disease (OR = 1.43), and all-cause death (OR = 1.41), while other disease events showed no change.

**Table 2.** Mean alcohol consumption in equivalent glasses of wine per week by group and ADH1B status.

| Group | Wildtype,<br>glasses/week | Variant,<br>glasses/week | Increase for wild type,<br>glasses/week |
| --- | --- | --- | --- |
| all | 8.2 | 6.1 | 2.1 |
| male | 11.2 | 8.4 | 2.8 |
| female | 5.5 | 3.9 | 1.6 |
| current smoker | 11.1 | 7.6 | 3.5 |
| never-smoker | 6.5 | 4.6 | 1.9 |

Drinking varied by sex (men: 11.1 glasses/week vs. women: 5.4 glasses/week) and by smoking status (current smokers: 9.8 glasses/week vs. never-smokers: 6.5 glasses/week) in stratified analyses (Table 2). One additional drink per day corresponded to higher systolic blood pressure, BMI, waist circumference, body fat percentage, and LDL in all groups. Men additionally had higher risk of heart failure (OR=1.76), atrial fibrillation (OR=1.35), and type 2 diabetes (OR=1.31) with one additional drink per day, while there was no difference in risk for these diseases in women. Women did not have any increase in myocardial infarction with alcohol use. We measured statistical evidence for heterogeneity of the OR between men and women for myocardial infarction (p=0.070), heart failure (p=0.048), atrial fibrillation (p=0.041), type 2 diabetes (p=0.068), all CVD (p=0.013), and all-cause death (p=0.015).

Sensitivity analyses in subgroups not taking medications for diabetes, hypertension, and lipids showed negligible differences in results for HbA1C, systolic/diastolic blood pressure, and LDL/triglycerides, respectively (S5.2). Analysis of blood count data by instrument variable showed that those with the wildtype, who drink more, had a lower hemoglobin (−0.02 g/dL, p=0.0018) and hematocrit (−0.09%, p=0.0024), and a higher mean corpuscular volume (+0.27 fL, p=2.4 x 10^−13^) (Table S5.3).

In summary statistics from external datasets, alcohol was predictive of stroke and ischemic stroke (MEGASTROKE), type 2 diabetes (DIAMANTE), BMI (GIANT-UK Biobank metaanalysis), and waist circumference (EXTEND) (Figure 1).

## Discussion

Our principal analysis within the UK Biobank suggests that alcohol is causally associated not only with a range of adverse cardiovascular related outcomes such as myocardial infarction, stroke, and all-cause death, but also multiple traditional risk factors for these outcomes that likely mediate these observed effects including hypertension (systolic blood pressure), obesity (waist circumference, body fat percentage), and atherogenic dyslipidemia (cholesterol, LDL). Our findings in the UK Biobank were further supported by analyses involving external datasets for a subset of these outcomes including obesity (BMI, waist circumference), stroke, and Type 2 diabetes. Our MR analyses gave discrepant results when compared to the analogous observational analysis for these phenotypes, which suggests the presence of residual confounding from unmeasured factors in observational analysis.

Two findings in our analyses are inconsistent with the hypothesis of a causal negative effect on cardiometabolic outcomes mediated through risk factors. First, we found alcohol to be causally associated with a lower level of triglycerides in the UK Biobank, which would be expected to reduce the risk of atherosclerosis related outcomes. We note this finding is contrary to what has been observed experimentally [26]; but consistent with other MR studies [27–29]. Although multiple MR studies suggest triglycerides are causally associated with CVD, clinical trials of triglyceride-lowering agents have not consistently supported this relationship leading to the conclusion that how triglycerides are lowered plays an important role in whether that lowering translates to cardiovascular benefit. Second, we found alcohol to be inversely associated with HbA1C which would be expected to reduce the risk of Type 2 diabetes. We suspect this counterintuitive association observed may be a technical artifact driven by the presence of a mild (possibly nutritional) macrocytosis we observed among participants not carrying the minor allele at rs1229984 that biases HgA1C levels downwards without truly altering the risk of diabetes[30, 31].

Our stratified analyses suggest that alcohol is more harmful for men than for women for nearly all outcomes. We observed the largest differences in the point estimates of the harmful effect for heart-related outcomes (myocardial infarction, heart failure, and atrial fibrillation) with a smaller difference observed for type 2 diabetes. For most outcomes, the point estimate of effect of alcohol for women was near 1. Tests for heterogeneity of odds ratios for outcomes which appeared to differ between men and women were not definitive as to whether a true difference exists but strongly suggestive. Larger samples sizes are needed to more reliably document a statistically significant modification of effect of alcohol between men and women. If confirmed, these findings would suggest that causal negative effects of alcohol may only begin to express themselves at a higher consumption level despite differences in body surface area and rate of metabolism between females and males. We did not observe the same trend between our smoking subgroup analysis where the effect of drinking did not differ substantially between smokers and non-smokers for most outcomes.

Our findings are largely consistent with the existing literature of MR studies of alcohol and cardiovascular risk factors and outcomes which have found alcohol to be harmful for CVD related outcomes and most cardiometabolic risk factors [27–29, 32, 33]. For example, the first large MR study of risk factors and outcomes using data gathered from over 56 cohort studies of individuals of European ancestry and the same genetic instrument in *ADH1B* found moderate alcohol use to be associated with higher systolic blood pressure, waist circumference, BMI, LDL, and risk of coronary heart disease [28]. Another smaller MR study in Danes also found a direct association with a higher BMI [27]. A study of the China Kadoorie biobank (>500,000) used a combination of instruments in both *ALDH2* and *ADH1B* found alcohol to be positively associated with increasing systolic blood pressure, HDL, ischemic stroke, and intracerebral hemorrhage, but no effect was found for myocardial infarction [32]. Another recent study of alcohol and CVD outcomes in the UK Biobank using a multi-SNP instrument variable found an increase in blood pressure, stroke, and peripheral artery disease [29]. The inverse association of alcohol with triglycerides that we observed has also been shown in several other MR studies[27–29]. Diabetes has previously been found to have no association with alcohol in an MR meta-analysis (>14,000 cases) as instrumented by the same variant we used [28]. However, another study in a Chinese population using the *ALDH2* rs671 variant, which has a more profound effect on alcohol intake, found a higher risk of diabetes with increasing alcohol [33]. Unique contributions of our study include the identification of the harmful effects of alcohol on body fat percentage and all-cause death. Our study also adds to the existing literature by showing sex-stratified differences which merit further investigation.

Major strengths of our study include the use of a single SNP as a genetic instrument to predict causality of alcohol consumption combined with a large sample size. Using a single instrument within an alcohol metabolizing gene that is strongly associated with the exposure maximizes the probability that the all assumptions of an MR study have been met and the results are accurately reflecting a relationship that is free from any residual confounding [34]. A potential weakness may be the generalizability of our study given the well-established healthy cohort effect observed for the UK Biobank [35], although the healthy cohort may have helped by minimizing the inclusion of subjects with alcohol use disorder and/or moderate but still high-risk use of alcohol (e.g. binge drinkers). Our results are also limited to UK residents and therefore may vary somewhat in other populations although MR studies to date in other populations including Chinese are largely consistent with our findings[27, 28, 32, 33]. More research is needed on the determination of the causal effects of alcohol consumption in race/ethnic groups other than Europeans and East Asians to determine if effects observed to date generalize to all major race/ethnic groups.

Proposed mechanisms for the negative effect of alcohol on cardiometabolic disease include a pathway via raised blood pressure [36] and atherogenic lipids [37] as well as increased adiposity and subsequent risk of Type2 diabetes, consistent with our MR findings and those of others. Raised HDL has been proposed as a protective factor but our MR results do not support that such elevations are directly related to alcohol among a population of predominantly moderate alcohol users. The same relationship with HDL has been observed in other MR studies[28, 29, 33]. Additionally, multiple lines of evidence now suggest that HDL levels are not causally associated with heart disease [17, 18] but instead serve as a marker of a variety of factors that may or may not affect the risk of CVD. In this context, one can speculate that physical activity raises HDL in a health-promoting way [38], while alcohol consumption does not. Further harm of alcohol for stroke could come from alcohol induced thrombocytopenia (hemorrhagic stroke) and reduced fibrinolysis (ischemic stroke) with alcohol use [39].

In conclusion, our analysis adds to the mounting evidence using MR that alcohol use does not improve cardiovascular health even in moderate amounts and likely worsens it when all other factors are considered. Given this evidence and the fact that alcohol is implicated in a number of public health concerns not directly related to cardiometabolic health, including addictive disorders, accidents, suicides, liver disease, and various types of cancers (e.g. esophageal, gastrointestinal, head and neck) [40], we believe it is time to reconsider current public health recommendations in the US and other countries which suggest that up to two drinks/day for men and one drink/day for women is not harmful and possibly beneficial to cardiovascular health [41]. This reconsideration is also supported by a more recent observational study that considered the full spectrum of alcohol related health consequences across the entire age spectrum and concluded that the level of consumption that minimizes health loss is zero [40]. Properly conducted randomized control trials [42] may one day more reliably inform us on this matter but, until that time comes, Mendelian randomization analyses provides an acceptable alternative to help inform health policy in this respect.

## Tables and figures

**Figure 1**

See separate PDFs

Caption title: Results from Mendelian randomization of outcome variables in UK Biobank and external datasets

Caption: Significant results shown in black with circles; non-significant results shown in gray with squares.

**Figure 1a:**
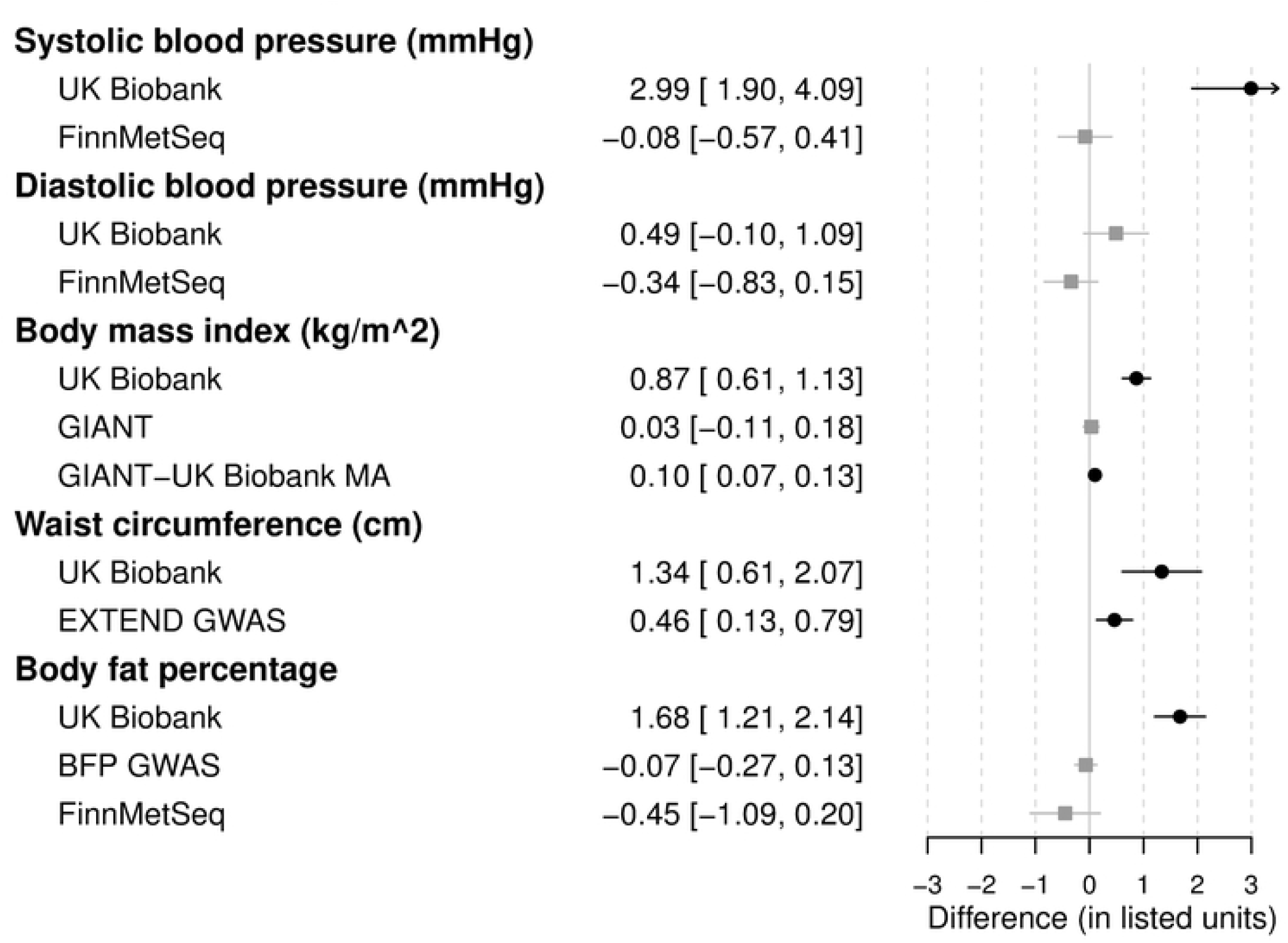
Mendelian randomization results for blood pressure and anthropometric measures

**Figure 1b:**
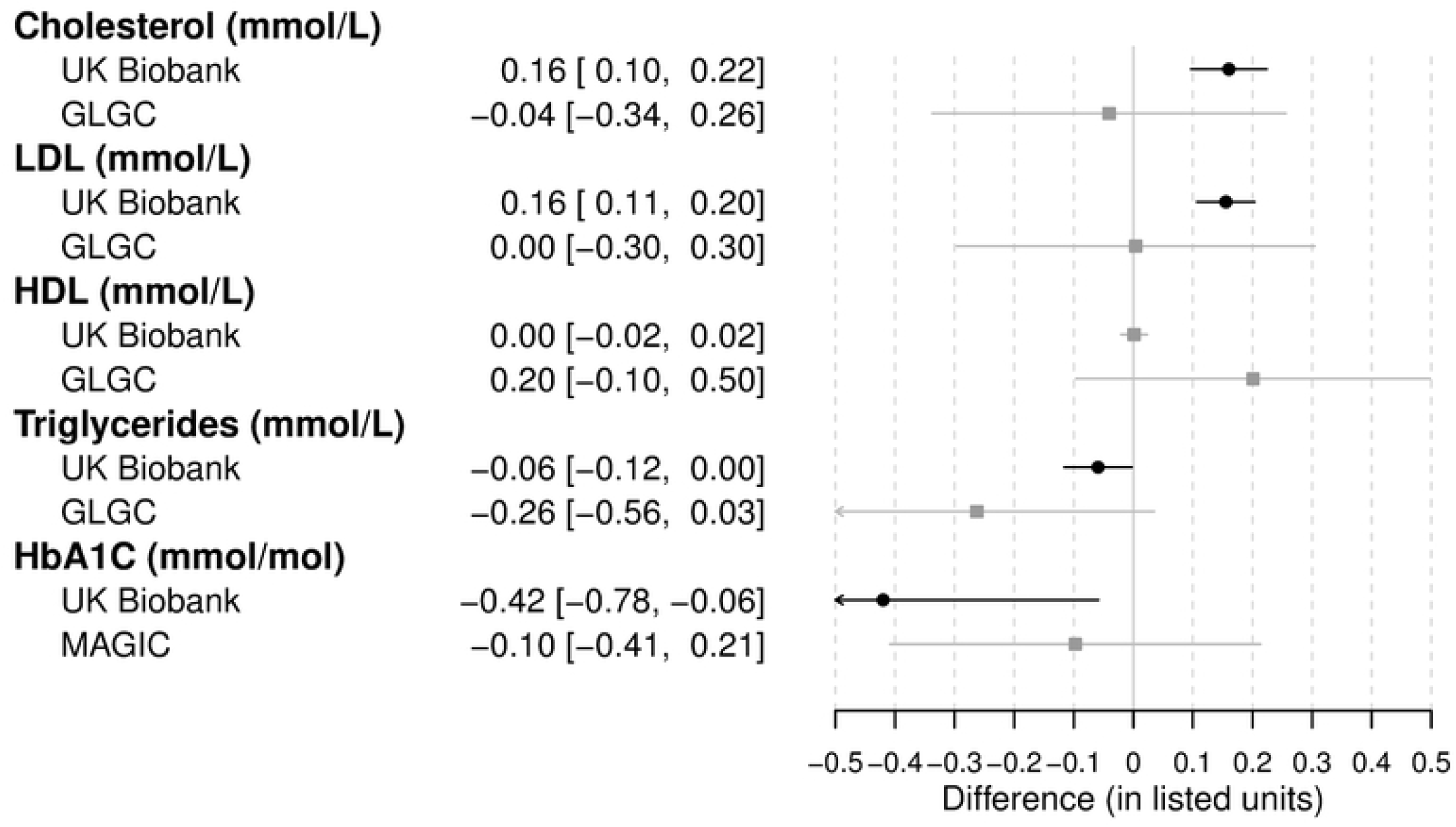
Mendelian randomization results for biochemistry variables

**Figure 1c:**
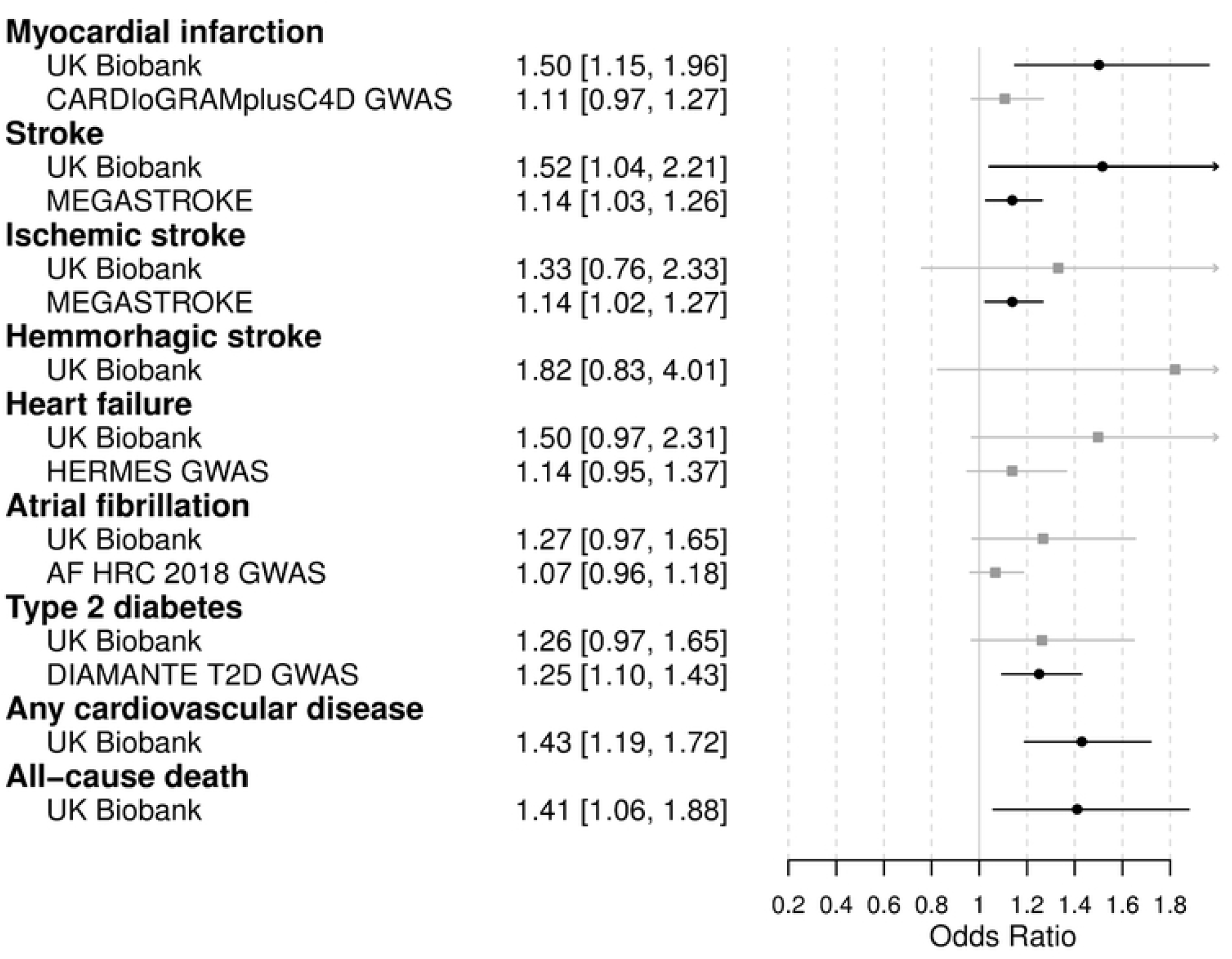
Mendelian randomization results for disease outcomes and death

**Figure 2**

See separate PDFs

Caption title: Results from Mendelian randomization of outcome variables in UK Biobank stratified by sex and smoking status

Caption: Significant results shown in black with circles; non-significant results shown in gray with squares.

**Figure 2a:**
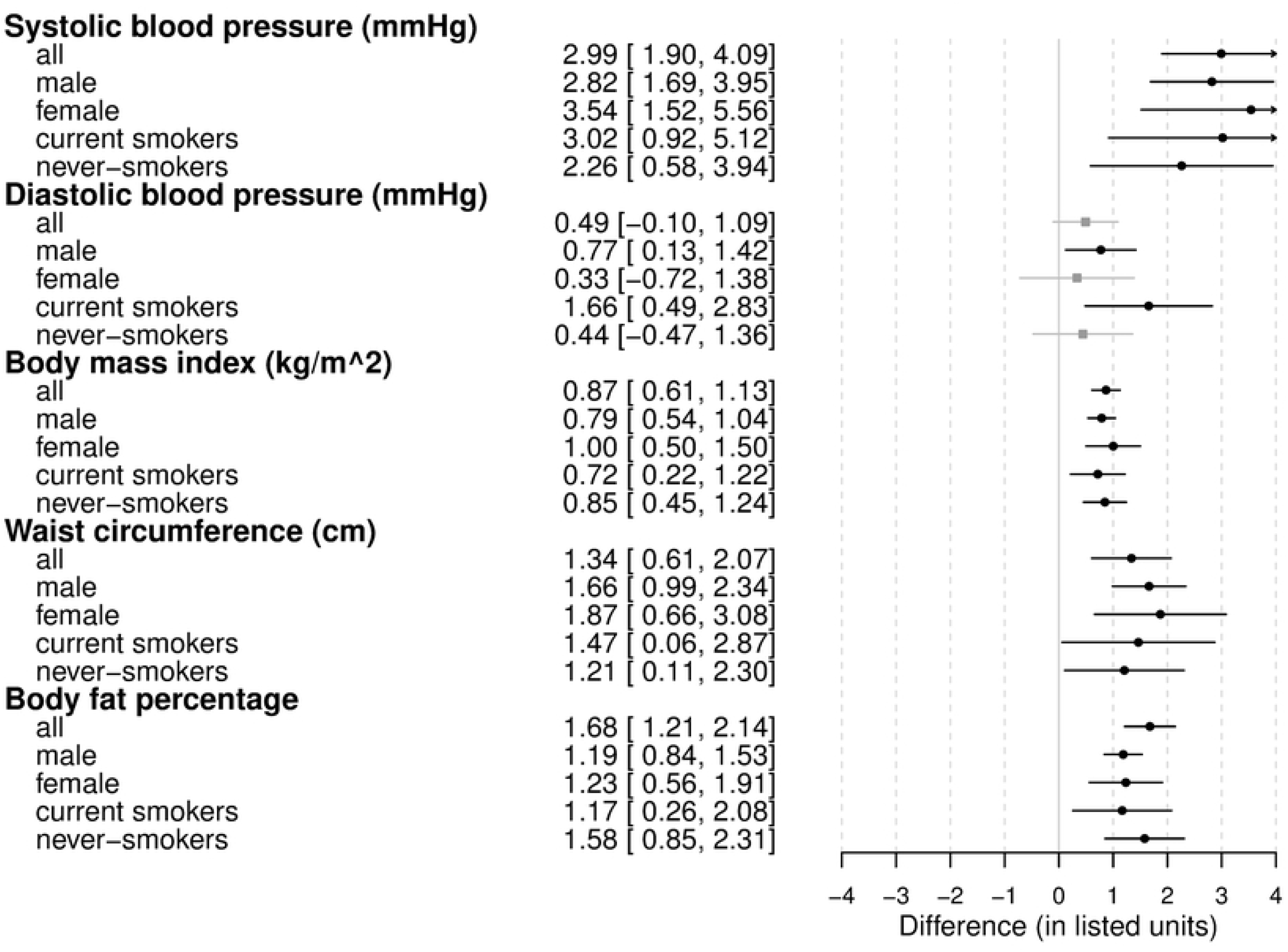
Stratified Mendelian randomization results for blood pressure and anthropometric measures

**Figure 2b:**
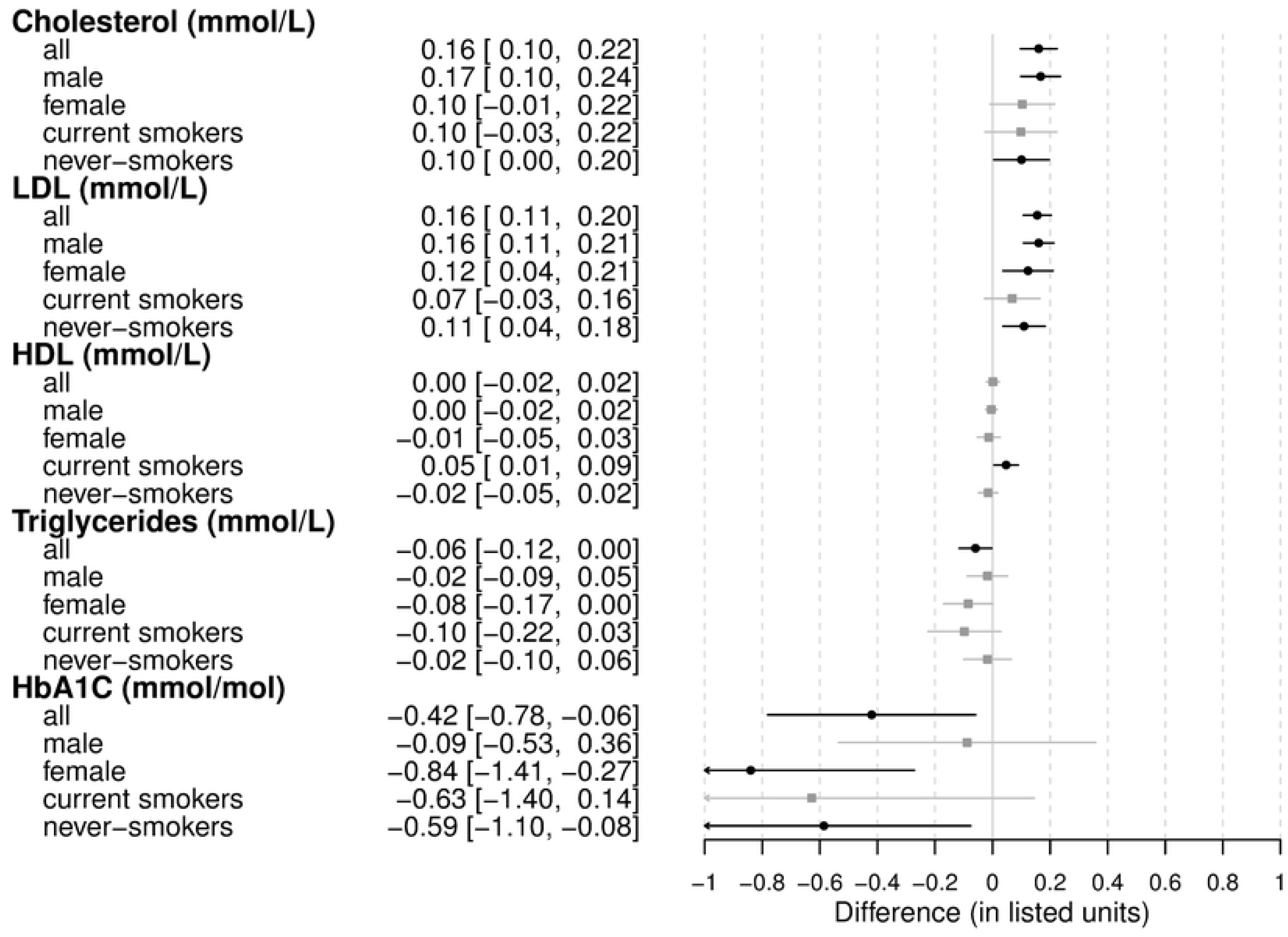
Stratified Mendelian randomization results for biochemistry variables

**Figure 2c:**
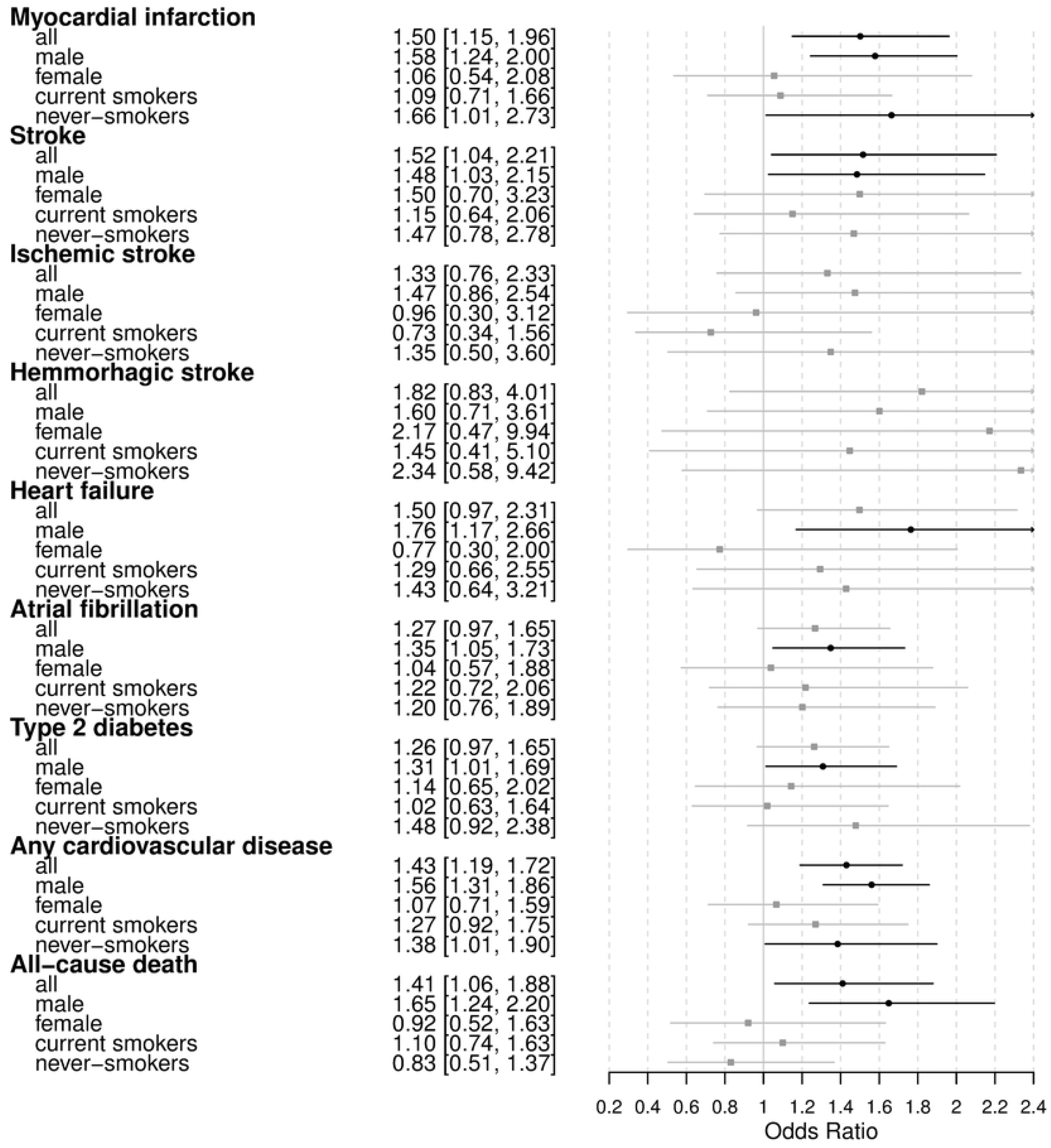
Stratified Mendelian randomization results for disease outcomes and death

## Supplement

### Supplemental methods detail

#### S1. Data transformations

##### S1.1 Disease events

Disease events were defined according to each of the following ICD codes and extracted from hospital data (using both primary and secondary diagnoses).

| Event | ICD-9 | ICD-10 | OPCS-4 |
| --- | --- | --- | --- |
| Myocardial infarction | 410*, 411* | I20.0, I21*, I22* | K40*, K41*, K42*, K43*, K44*, K45*, K46*, K49*, K50*, K75* |
| All stroke | 430*, 431*, 432*, 433*, 434*, 435*, 436* | I60*, I61*, I62*, I63*, G45*, I64* | n/a |
| Ischemic stroke | 433*, 434* | I63* | n/a |
| Hemorrhagic stroke | 430*, 431*, 432* | I60*, I61*, I62* | n/a |
| Heart failure | 428* | I50* | n/a |
| Atrial fibrillation | 427.3* | I48* | K52.1, K62.2, K62.3, K62.4 |

\* Including all subcategories

##### S1.2 Diabetes

Diabetes was defined using self-reported diabetes (question # 1221, 1222, 1223), self-reported medical conditions (question # 20002), self-reported medications (question # 20003), HbA1C results (question # 30750), age of diagnosis (question # 2976), started insulin within one year of diagnosis (question # 2986), diabetes diagnosed by a doctor (question # 2443), diabetes gestational only (question # 4041).

We flagged individuals who had suspected Type 1 diabetes. We considered individuals to have Type 2 diabetes if they were not flagged for Type 1, and any of the following was true:

- HbA1C exceeded threshold of 48 mmol/mol
- Type 2 diabetes was in their self-reported medical conditions
- Diabetes medications were self-reported
- Diabetes had been diagnosed by a doctor but was not gestational only

We flagged individuals for suspected type 1 diabetes if:

- Type 1 diabetes was in reported conditions, OR
- Diabetes was diagnosed before age 35, OR
- Insulin began within a year of diagnosis.

#### S2. Observational analysis

In the partially-adjusted model, we used the Townsend index to adjust for socioeconomic status. Ethnicity was grouped by the highest tree-structure group of the 21000 variable. Region was encoded as England, Scotland, or Wales. Antihypertensive and lipid-lowering medications were extracted from variables 6153 and 6177. Diabetes medications came from variables 6153, 6177, and 20003. Sex, age, and smoking status also came from survey data. Fasting time (variable 74) was also used in all biomarker outcome models.

Strikethrough text indicates a variable was not included in the fully-adjusted model.

| Outcome | Fully adjusted includes all the partially adjusted plus: |
| --- | --- |
| BMI, waist circumference, body fat percentage | HbA1c, diabetes medications, ~~BMI, waist circumference,~~ SBP, DBP, anti-hypertensive, LDL, HDL, triglycerides, lipid-lowering |
| SBP and DBP | HbA1c, diabetes medications, BMI, waist circumference, ~~SBP, DBP,~~ anti-hypertensive, LDL, HDL, triglycerides, lipid-lowering |
| Type 2 diabetes | ~~HbA1c, diabetes medications,~~ BMI, waist circumference, SBP, DBP, anti-hypertensive, LDL, HDL, triglycerides, lipid-lowering |
| Cholesterol, LDL, HDL, triglycerides | HbA1c, diabetes medications, BMI, waist circumference, SBP, DBP, anti-hypertensive, ~~LDL, HDL, triglycerides,~~ lipid-lowering |
| HBA1C | ~~HbA1c,~~ diabetes medications, BMI, waist circumference, SBP, DBP, anti-hypertensive, LDL, HDL, triglycerides, lipid-lowering |
| Heart disease events (CHD, stroke, etc) | HbA1c, diabetes medications, BMI, waist circumference, SBP, DBP, anti-hypertensive, LDL, HDL, triglycerides, lipid-lowering |

#### S3. Mendelian randomization analysis

Where possible, we utilized the Cardiovascular Disease, Cerebrovascular Disease, and Type 2 Diabetes Knowledge Portals.

**Table S3:** Data sources for Mendelian randomization. * indicates the source also includes UK Biobank data. CVDKP = Cardiovascular disease knowledge portal [43], http://www.broadcvdi.org/home/portalHome CDKP = Cerebrovascular disease knowledge portal [44], http://cerebrovascularportal.org/ T2DKP = Type 2 diabetes knowledge portal [45], http://www.type2diabetesgenetics.org/

| Name | Extracted summary statistics for phenotypes | Population ancestry | n | From |
| --- | --- | --- | --- | --- |
| CARDIoGRAMplusC4D with 1000 genomes GWAS meta-analysis [46] | Myocardial infarction | Mixed | 184,305 | http://www.cardiogramplusc4d.org/data-downloads/ |
| MEGASTROKE [47] | All stroke, Ischemic stroke | Mixed | 521,612 | CDKP |
| HERMES Heart Failure GWAS* [48] | Heart failure | European | 972,032 | CVDKP |
| 2018 AF HRC GWAS [49] | Atrial fibrillation | Mixed | 588,190 | CVDKP |
| DIAMANTE T2D GWAS* [50] | Type 2 diabetes | European | 898,130 | T2DKP |
| FinnMetSeq exome sequence analysis [51] | Systolic blood pressure, diastolic blood pressure, body fat percentage | European | 19,291 | T2DKP |
| MAGIC GWAS [52] | HbA1C | European | 5,318 | T2DKP |
| GIANT 2018 BMI, Height exome chip analysis [53] | BMI | European | 449,889 | https://portals.broadinstitute.org/collaboration/giant/index.php/GIANT_consortium_data_files |
| GIANT-UK Biobank GWAS Meta-analysis* [54] | BMI | European | 694,649 | T2DKP |
| EXTEND GWAS [55] | Waist circumference | European | 7,159 | T2DKP |
| Body fat percentage GWAS [56] | Body fat percentage | Mixed (89,267 European) | 100,716 | T2DKP |
| GLGC exome chip analysis [57] | Total cholesterol, LDL, HDL, triglycerides | Mixed (273,050 European) | ~300,000 | https://my.locuszoom.org/gwas/919709/ |

### Supplemental results

#### S4. Observational analysis results

All results are compared to a reference of no alcohol consumption

SBP: Systolic blood pressure (mmHg)

DBP: Diastolic blood pressure (mmHg)

BMI: Body mass index (kg/m^2)

WAIST: Waist circumference (cm)

BFP: Body fat percentage

CHOL: Cholesterol (mmol/L)

LDL: Low-density lipoprotein (mmol/L)

HDL: High-density lipoprotein (mmol/L)

TG: Triglycerides (mmol/L)

HBA1C: Glycated hemoglobin (mmol/mol)

T2D: type 2 diabetes

MI: myocardial infarction

ALLSTROKE: all types of stroke

ISTROKE: ischemic stroke

HSTROKE: hemorrhagic stroke

HF: heart failure

AFIB: atrial fibrillation

CVD: any cardiovascular disease (MI, ALLSTROKE, ISTROKE, HSTROKE, HF, AFIB) Death: all-cause death

Figure S4.1: Observational analysis results for effect size for given drinking category (units listed with abbreviations) for fully adjusted model.

Figure S4.2: Observational analysis results for hazard ratio (event outcomes) or odds ratio (type 2 diabetes) for fully adjusted model.

#### S4.3 Betas and confidence intervals for continuous measures, basic model. Reference: no alcohol consumption

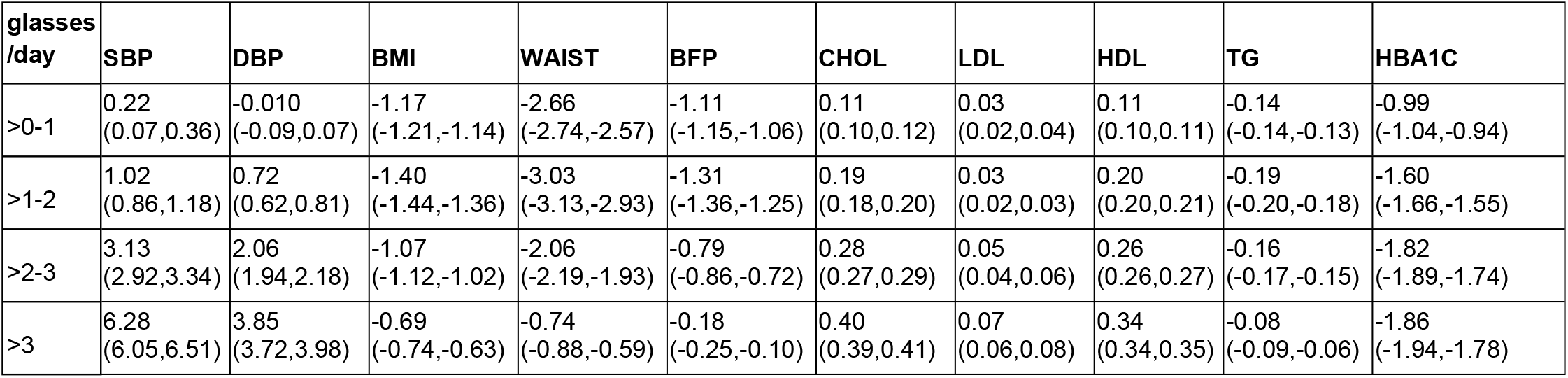

| **glasses**<br>**/day** | **SBP** | **DBP** | **BMI** | **WAIST** | **BFP** | **CHOL** | **LDL** | **HDL** | **TG** | **HBA1C** |
| --- | --- | --- | --- | --- | --- | --- | --- | --- | --- | --- |
| >0-1 | 0.22<br>(0.07,0.36) | -0.010<br>(-0.09,0.07) | -1.17<br>(-1.21,-1.14) | -2.66<br>(-2.74,-2.57) | -1.11<br>(-1.15,-1.06) | 0.11<br>(0.10,0.12) | 0.03<br>(0.02,0.04) | 0.11<br>(0.10,0.11) | -0.14<br>(-0.14,-0.13) | -0.99<br>(-1.04,-0.94) |
| >1-2 | 1.02<br>(0.86,1.18) | 0.72<br>(0.62,0.81) | -1.40<br>(-1.44,-1.36) | -3.03<br>(-3.13,-2.93) | -1.31<br>(-1.36,-1.25) | 0.19<br>(0.18,0.20) | 0.03<br>(0.02,0.03) | 0.20<br>(0.20,0.21) | -0.19<br>(-0.20,-0.18) | -1.60<br>(-1.66,-1.55) |
| >2-3 | 3.13<br>(2.92,3.34) | 2.06<br>(1.94,2.18) | -1.07<br>(-1.12,-1.02) | -2.06<br>(-2.19,-1.93) | -0.79<br>(-0.86,-0.72) | 0.28<br>(0.27,0.29) | 0.05<br>(0.04,0.06) | 0.26<br>(0.26,0.27) | -0.16<br>(-0.17,-0.15) | -1.82<br>(-1.89,-1.74) |
| >3 | 6.28<br>(6.05,6.51) | 3.85<br>(3.72,3.98) | -0.69<br>(-0.74,-0.63) | -0.74<br>(-0.88,-0.59) | -0.18<br>(-0.25,-0.10) | 0.40<br>(0.39,0.41) | 0.07<br>(0.06,0.08) | 0.34<br>(0.34,0.35) | -0.08<br>(-0.09,-0.06) | -1.86<br>(-1.94,-1.78) |

##### S4.4 Betas and confidence intervals for continuous measures, fully-adjusted model. Reference: no alcohol consumption

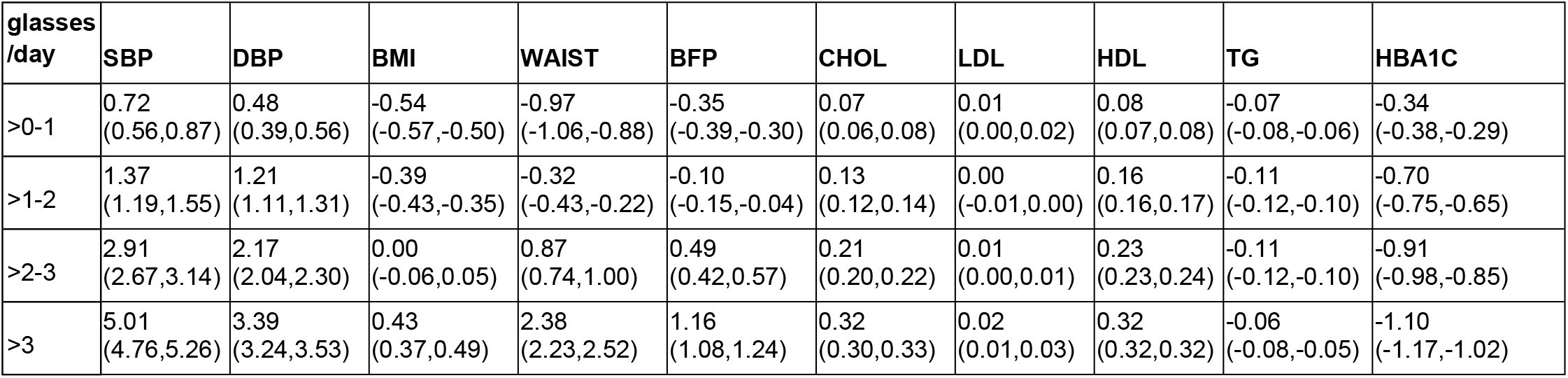

*S4.4 Betas and confidence intervals for continuous measures, fully-adjusted model. Reference: no alcohol consumption.*
| **glasses**<br>**/day** | **SBP** | **DBP** | **BMI** | **WAIST** | **BFP** | **CHOL** | **LDL** | **HDL** | **TG** | **HBA1C** |
| --- | --- | --- | --- | --- | --- | --- | --- | --- | --- | --- |
| >0-1 | 0.72<br>(0.56,0.87) | 0.48<br>(0.39,0.56) | -0.54<br>(-0.57,-0.50) | -0.97<br>(-1.06,-0.88) | -0.35<br>(-0.39,-0.30) | 0.07<br>(0.06,0.08) | 0.01<br>(0.00,0.02) | 0.08<br>(0.07,0.08) | -0.07<br>(-0.08,-0.06) | -0.34<br>(-0.38,-0.29) |
| >1-2 | 1.37<br>(1.19,1.55) | 1.21<br>(1.11,1.31) | -0.39<br>(-0.43,-0.35) | -0.32<br>(-0.43,-0.22) | -0.10<br>(-0.15,-0.04) | 0.13<br>(0.12,0.14) | 0.00<br>(-0.01,0.00) | 0.16<br>(0.16,0.17) | -0.11<br>(-0.12,-0.10) | -0.70<br>(-0.75,-0.65) |
| >2-3 | 2.91<br>(2.67,3.14) | 2.17<br>(2.04,2.30) | 0.00<br>(-0.06,0.05) | 0.87<br>(0.74,1.00) | 0.49<br>(0.42,0.57) | 0.21<br>(0.20,0.22) | 0.01<br>(0.00,0.01) | 0.23<br>(0.23,0.24) | -0.11<br>(-0.12,-0.10) | -0.91<br>(-0.98,-0.85) |
| >3 | 5.01<br>(4.76,5.26) | 3.39<br>(3.24,3.53) | 0.43<br>(0.37,0.49) | 2.38<br>(2.23,2.52) | 1.16<br>(1.08,1.24) | 0.32<br>(0.30,0.33) | 0.02<br>(0.01,0.03) | 0.32<br>(0.32,0.32) | -0.06<br>(-0.08,-0.05) | -1.10<br>(-1.17,-1.02) |

##### S4.5 Odds ratios and confidence intervals for type 2 diabetes, basic model. Reference: no alcohol consumption

*S4.5 Odds ratios and confidence intervals for type 2 diabetes, basic model. Reference: no alcohol consumption.*
| **glasses/day** | **T2D** |
| --- | --- |
| >0-1 | 0.59<br>(0.57,0.61) |
| >1-2 | 0.44<br>(0.42,0.46) |
| >2-3 | 0.43 (0.4,0.45) |
| >3 | 0.47 (0.45,0.5) |

##### S4.6 Odds ratios and confidence intervals for type 2 diabetes, fully-adjusted model. Reference: no alcohol consumption

| **glasses/day** | **T2D** |
| --- | --- |
| >0-1 | 0.81<br>(0.77,0.84) |
| >1-2 | 0.69<br>(0.65,0.73) |
| >2-3 | 0.68<br>(0.63,0.73) |
| >3 | 0.75<br>(0.7,0.81) |

##### S4.7 Hazard ratios and confidence intervals for binary measures (diseases and events), basic model. Reference: no alcohol consumption

| **glasses/day** | **MI** | **ALLSTROKE** | **ISTROKE** | **HSTROKE** | **HF** | **AFIB** | **CVD** | **Death** |
| --- | --- | --- | --- | --- | --- | --- | --- | --- |
| >0-1 | 0.80<br>(0.76,0.84) | 0.89<br>(0.84,0.95) | 0.85<br>(0.78,0.93) | 0.93<br>(0.82,1.05) | 0.69<br>(0.64,0.74) | 0.88<br>(0.84,0.92) | 0.83<br>(0.81,0.86) | 0.77<br>(0.74,0.80) |
| >1-2 | 0.70<br>(0.66,0.74) | 0.83<br>(0.78,0.89) | 0.86<br>(0.78,0.95) | 0.79<br>(0.68,0.92) | 0.61<br>(0.56,0.66) | 0.89<br>(0.85,0.94) | 0.78<br>(0.75,0.80) | 0.71<br>(0.68,0.74) |
| >2-3 | 0.68<br>(0.63,0.72) | 0.79<br>(0.72,0.87) | 0.75<br>(0.65,0.86) | 0.93<br>(0.77,1.12) | 0.63<br>(0.57,0.70) | 0.94<br>(0.88,0.99) | 0.79<br>(0.75,0.82) | 0.71<br>(0.67,0.75) |
| >3 | 0.69<br>(0.65,0.74) | 0.98<br>(0.90,1.08) | 1.00<br>(0.88,1.14) | 1.07<br>(0.88,1.29) | 0.75<br>(0.68,0.83) | 1.10<br>(1.04,1.17) | 0.88<br>(0.85,0.92) | 0.91<br>(0.87,0.96) |

##### S4.8 Hazard ratios and confidence intervals for binary measures (diseases and events), fully-adjusted model. Reference: no alcohol consumption

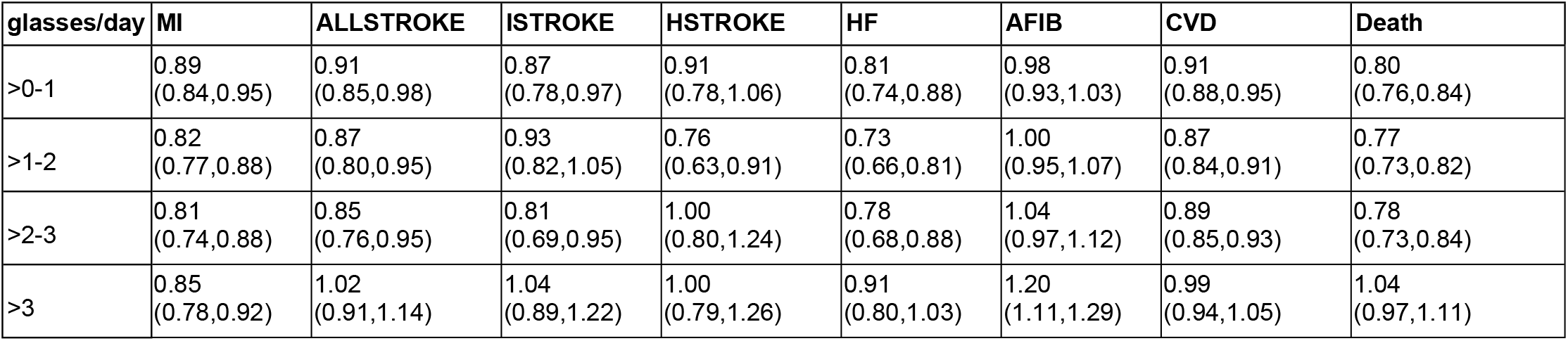

#### S5. Mendelian randomization results

##### S5.1 Characteristics of strata for the Mendelian randomization (MR) analysis

The total column is repeated from Table 1.

| Variable | MR - total | MR - male | MR - female | MR - nonsmokers | MR - smokers |
| --- | --- | --- | --- | --- | --- |
| n | 337484 | 156248 | 181236 | 183826 | 33977 |
| Sex |  |  |  |  |  |
| Female | 181236 (53.7) | 0 (0) | 181236 (100) | 107418 (58.4) | 15577 (45.8) |
| Male | 156248 (46.3) | 156248 (100) | 0 (0) | 76408 (41.6) | 18400 (54.2) |
| Age (years) | 67.6 (8) | 67.9 (8.1) | 67.4 (7.9) | 66.9 (8.1) | 65.9 (8.1) |
| Region |  |  |  |  |  |
| England | 297645 (88.2) | 138232 (88.5) | 159413 (88) | 161683 (88) | 29350 (86.4) |
| Wales | 14824 (4.4) | 6822 (4.4) | 8002 (4.4) | 8236 (4.5) | 1614 (4.8) |
| Scotland | 25015 (7.4) | 11194 (7.2) | 13821 (7.6) | 13907 (7.6) | 3013 (8.9) |
| Townsend index | -1.6 (2.9) | -1.5 (3) | -1.6 (2.9) | -1.9 (2.7) | -0.1 (3.5) |
| Ethnic group |  |  |  |  |  |
| White | 337484 (100) | 156248 (100) | 181236 (100) | 183826 (100) | 33977 (100) |
| Asian or Asian British | 0 (0) | 0 (0) | 0 (0) | 0 (0) | 0 (0) |
| Black or Black British | 0 (0) | 0 (0) | 0 (0) | 0 (0) | 0 (0) |
| Mixed or other | 0 (0) | 0 (0) | 0 (0) | 0 (0) | 0 (0) |
| Don't know/refused | 0 (0) | 0 (0) | 0 (0) | 0 (0) | 0 (0) |
| Smoking status |  |  |  |  |  |
| Never-smoker | 183826 (54.5) | 76408 (48.9) | 107418 (59.3) | 183826 (100) | 0 (0) |
| Current smoker | 33977 (10.1) | 18400 (11.8) | 15577 (8.6) | 0 (0) | 33977 (100) |
| Former smoker | 118505 (35.1) | 60883 (39) | 57622 (31.8) | 0 (0) | 0 (0) |
| No response | 1176 (0.3) | 557 (0.4) | 619 (0.3) | 0 (0) | 0 (0) |
| Drinking status |  |  |  |  |  |
| Never-drinker | 10392 (3.1) | 2623 (1.7) | 7769 (4.3) | 8234 (4.5) | 674 (2) |
| Current drinker | 315257 (93.4) | 148479 (95) | 166778 (92) | 170381 (92.7) | 31430 (92.5) |
| Former drinker | 11543 (3.4) | 5017 (3.2) | 6526 (3.6) | 5084 (2.8) | 1806 (5.3) |
| No response | 292 (0.1) | 129 (0.1) | 163 (0.1) | 127 (0.1) | 67 (0.2) |
| Drinking frequency |  |  |  |  |  |
| Daily or almost daily | 72270 (21.4) | 41339 (26.5) | 30931 (17.1) | 30020 (16.3) | 9136 (26.9) |
| Three or four times a week | 81462 (24.1) | 42266 (27.1) | 39196 (21.6) | 44049 (24) | 6542 (19.3) |
| Once or twice a week | 88747 (26.3) | 40983 (26.2) | 47764 (26.4) | 52425 (28.5) | 7865 (23.1) |
| One to three times a month | 37367 (11.1) | 13662 (8.7) | 23705 (13.1) | 22816 (12.4) | 3720 (10.9) |
| Special occasions only | 35411 (10.5) | 10229 (6.5) | 25182 (13.9) | 21071 (11.5) | 4167 (12.3) |
| Never | 21991 (6.5) | 7656 (4.9) | 14335 (7.9) | 13354 (7.3) | 2491 (7.3) |
| No response | 236 (0.1) | 113 (0.1) | 123 (0.1) | 91 (0) | 56 (0.2) |
| Alcohol (weekly equivalent glasses of wine) | 8.1 (9.5) | 11.1 (11.2) | 5.4 (6.6) | 6.5 (7.8) | 10.9 (13.3) |
| Systolic blood pressure (mmHg) | 140.2 (19.7) | 143.2 (18.5) | 137.6 (20.3) | 139.7 (19.5) | 137.5 (19.5) |
| Diastolic blood pressure (mmHg) | 82.2 (10.7) | 84.1 (10.5) | 80.6 (10.5) | 82.2 (10.6) | 81.6 (10.9) |
| Body mass index (kg/m<sup>2</sup>) | 27.4 (4.7) | 27.8 (4.2) | 27 (5.1) | 27.1 (4.7) | 27.1 (4.8) |
| Waist circumference (cm) | 90.3 (13.5) | 97 (11.3) | 84.6 (12.5) | 88.7 (13.2) | 91.2 (13.5) |
| Body fat percentage | 31.4 (8.5) | 25.3 (5.8) | 36.6 (6.9) | 31.4 (8.6) | 29.9 (8.6) |
| Cholesterol (mmol/L) | 5.7 (1.1) | 5.5 (1.1) | 5.9 (1.1) | 5.7 (1.1) | 5.7 (1.2) |
| LDL (mmol/L) | 3.6 (0.9) | 3.5 (0.9) | 3.6 (0.9) | 3.6 (0.9) | 3.6 (0.9) |
| HDL (mmol/L) | 1.5 (0.4) | 1.3 (0.3) | 1.6 (0.4) | 1.5 (0.4) | 1.4 (0.4) |
| Triglycerides (mmol/L) | 1.8 (1) | 2 (1.1) | 1.6 (0.9) | 1.7 (1) | 2 (1.2) |
| HbA1c (mmol/mol) | 36 (6.5) | 36.3 (7.3) | 35.7 (5.7) | 35.5 (6) | 37.1 (7.2) |
| Glucose (mmol/L) | 5.1 (1.2) | 5.2 (1.4) | 5.1 (1) | 5.1 (1.1) | 5.1 (1.2) |
| Type 2 diabetes | 15493 (4.6) | 9879 (6.3) | 5614 (3.1) | 6402 (3.5) | 1715 (5) |
| Coronary heart disease | 16102 (4.8) | 12260 (7.8) | 3842 (2.1) | 6057 (3.3) | 2274 (6.7) |
| All stroke | 8044 (2.4) | 4825 (3.1) | 3219 (1.8) | 3498 (1.9) | 1193 (3.5) |
| Ischemic stroke | 3427 (1) | 2195 (1.4) | 1232 (0.7) | 1434 (0.8) | 561 (1.7) |
| Hemorrhagic stroke | 1877 (0.6) | 1002 (0.6) | 875 (0.5) | 811 (0.4) | 278 (0.8) |
| Heart failure | 5921 (1.8) | 4116 (2.6) | 1805 (1) | 2145 (1.2) | 920 (2.7) |
| Atrial fibrillation | 15483 (4.6) | 10425 (6.7) | 5058 (2.8) | 6766 (3.7) | 1530 (4.5) |
| Any cardiovascular disease | 35499 (10.5) | 24023 (15.4) | 11476 (6.3) | 14911 (8.1) | 4532 (13.3) |
| All-cause death | 13700 (4.1) | 8410 (5.4) | 5290 (2.9) | 5102 (2.8) | 2696 (7.9) |
| ADH1B status |  |  |  |  |  |
| Wildtype | 322519 (95.6) | 149158 (95.5) | 173361 (95.7) | 175669 (95.6) | 32518 (95.7) |
| Carrier | 14777 (4.4) | 7013 (4.5) | 7764 (4.3) | 8056 (4.4) | 1441 (4.2) |
| Homozygous minor allele | 188 (0.1) | 77 (0) | 111 (0.1) | 101 (0.1) | 18 (0.1) |

##### S5.2 Sensitivity analysis: Meds for diabetes, hypertension, and lipids

Our sensitivity analysis showed little difference in the overall outcome and no difference in the ultimate finding when excluding individuals with medications for diabetes, hypertension, or lowering of lipids.

| Group | HbA1C effect for +1 drink/day |
| --- | --- |
| Everyone (n=337,484) | -0.420 mmol/mol (-0.781, -0.059) |
| Diabetes medication takers excluded (n=325,466; 96%) | -0.412 mmol/mol (-0.684, -0.140) |

| Group | Systolic BP effect for +1 drink/day | Diastolic BP effect for +1 drink/day |
| --- | --- | --- |
| Everyone (n=337,484) | 2.99 mmHg (1.90, 4.09) | 0.492 mmHg (-0.102, 1.09) |
| Hypertensive medication takers excluded (n=267,379; 79%) | 2.77 mmHg (1.45, 4.09) | 0.573 mmHg (-0.149, 1.30) |

| Group | LDL effect for +1 drink/day | TG effect for +1 drink/day | HDL effect for +1 drink/day |
| --- | --- | --- | --- |
| Everyone (n=337,484) | 0.155 mmol/L (0.107, 0.204) | -0.059 mmol/L (-0.012, -0.002) | 0.0013 mmol/L (-0.021, 0.023) |
| Lipid lowering medication takers excluded (n=278,875; 83%) | 0.219 mmol/L (0.164, 0.273) | -0.080 mmol/L (-0.147, -0.013) | 0.0095 mmol/L (-0.017, 0.036) |

##### S5.3 Additional analysis of blood count data by instrument variable group

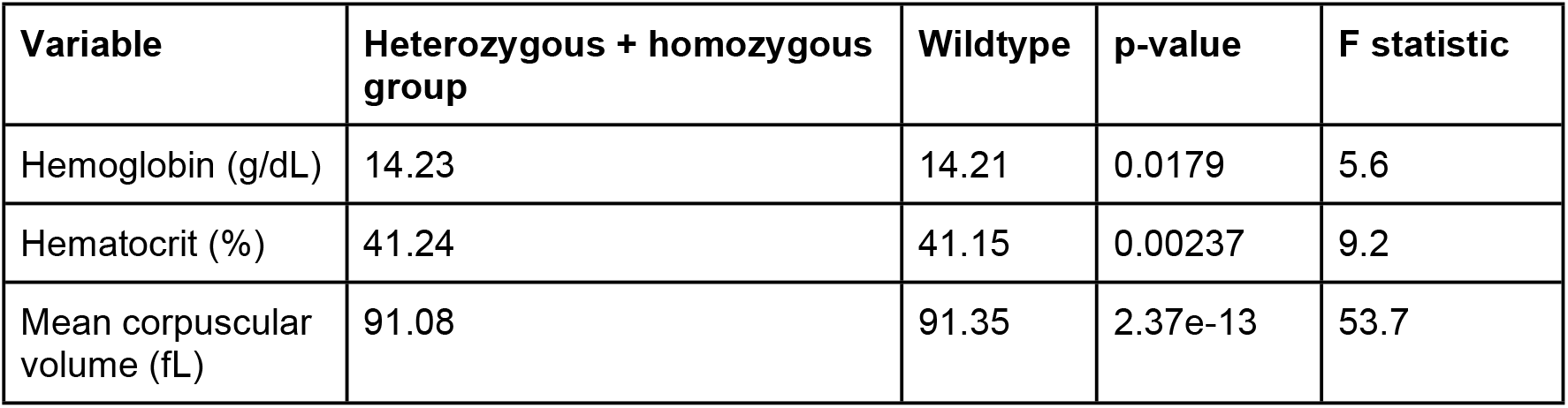

| Variable | Heterozygous + homozygous group | Wildtype | p-value | F statistic |
| --- | --- | --- | --- | --- |
| Hemoglobin (g/dL) | 14.23 | 14.21 | 0.0179 | 5.6 |
| Hematocrit (%) | 41.24 | 41.15 | 0.00237 | 9.2 |
| Mean corpuscular volume (fL) | 91.08 | 91.35 | 2.37e-13 | 53.7 |

---

**Figure S4-1.**
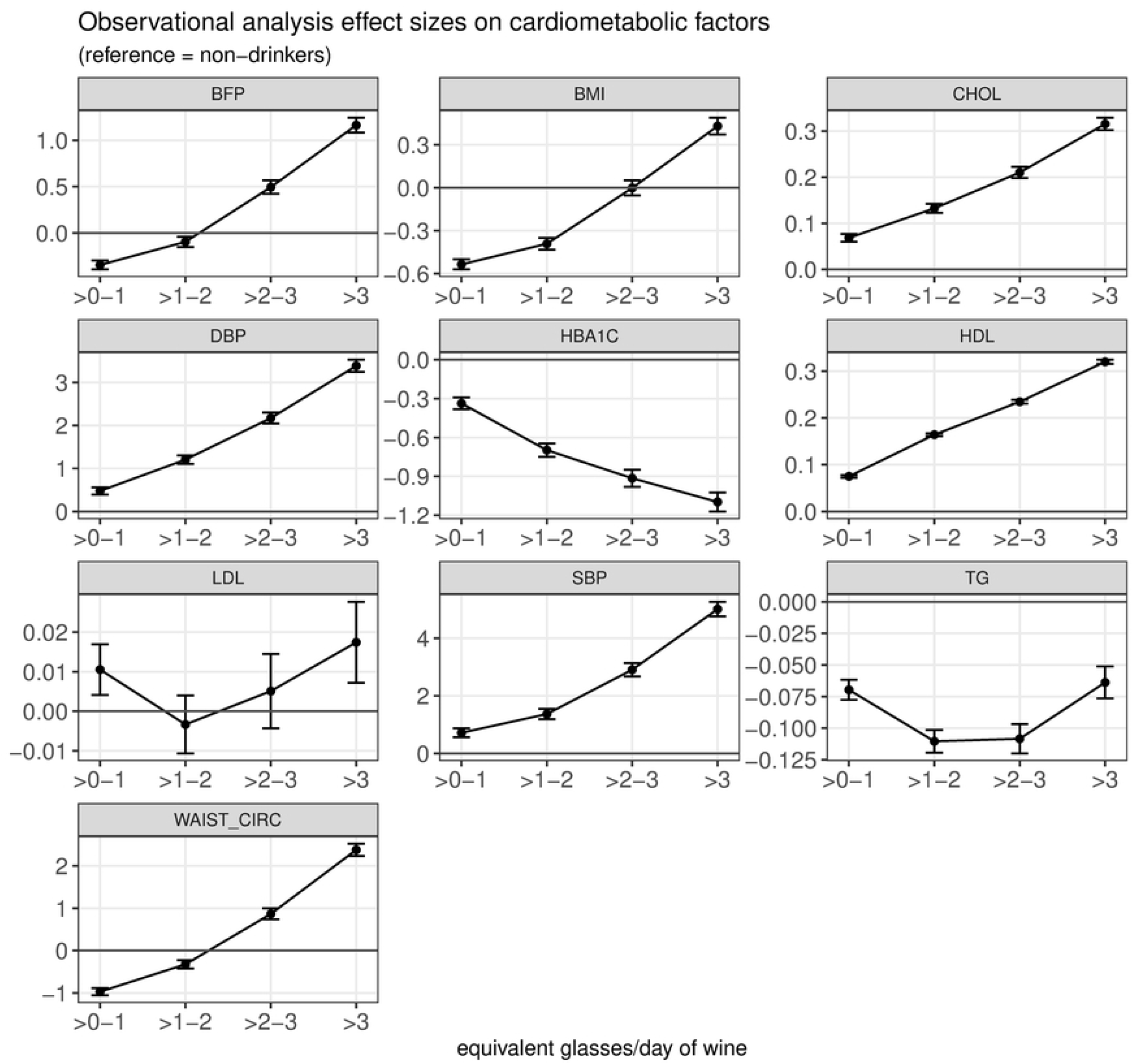

**Figure S4-2.**
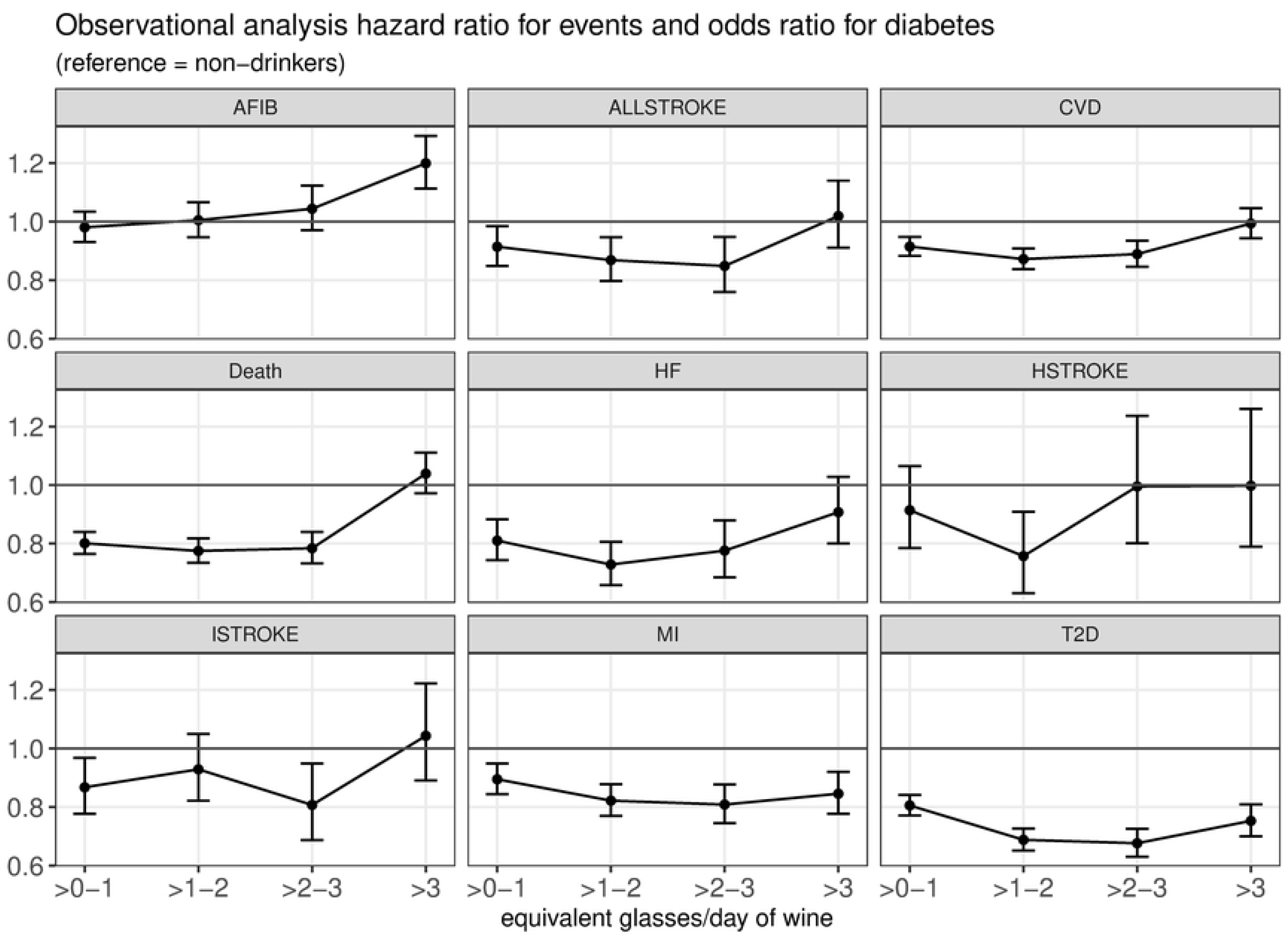

